## Supplementary Figures for "A strategy to incorporate prior knowledge into correlation network cutoff selection"

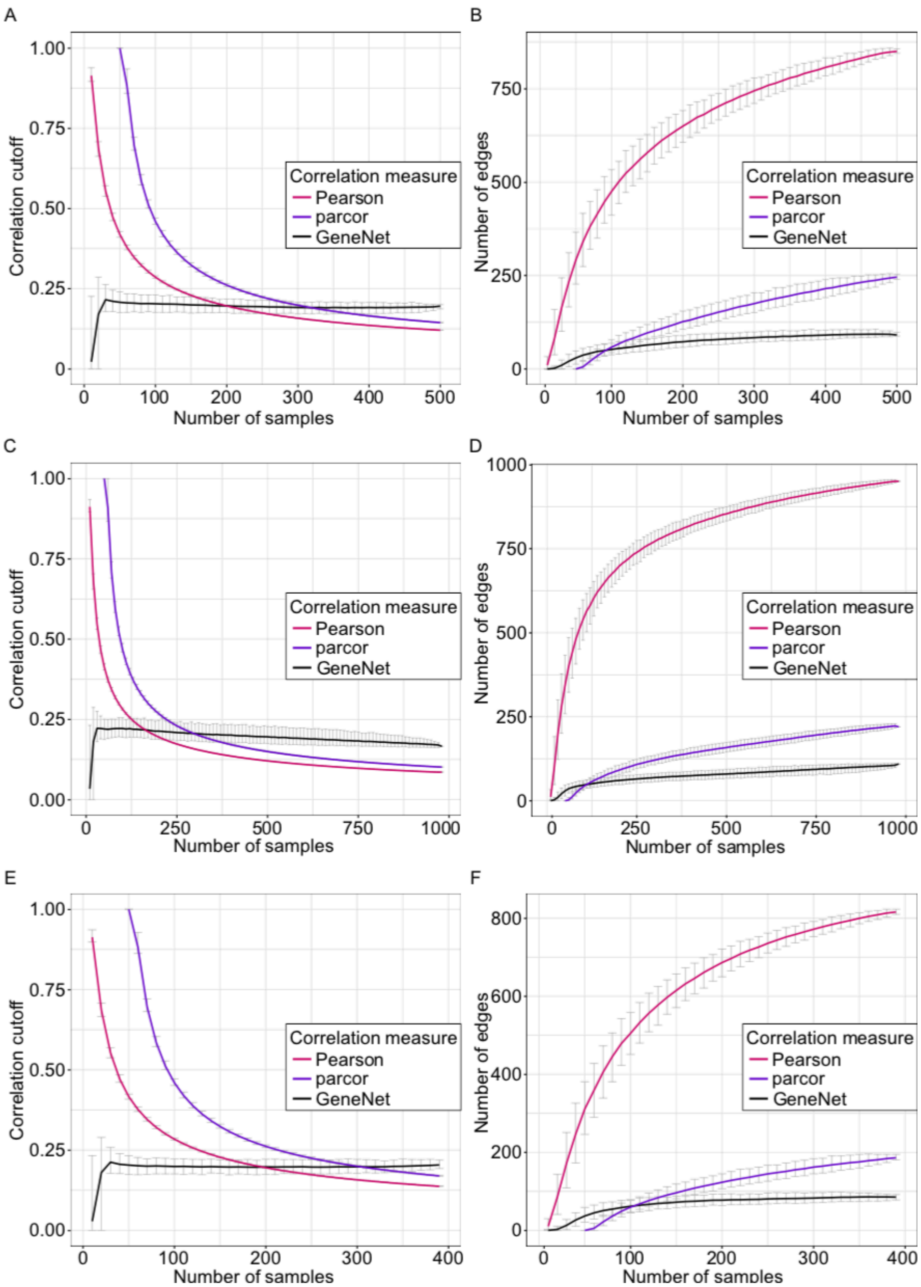


**Figure S1: Size dependence of statistical cutoffs in the glycomics replication cohorts.**

**A, C, E** Correlation cutoff (0.01 FDR, Benjamini-Hochberg) as a function of the dataset sample size for the three correlation measures considered in the Korčula 2010, Split and Vis cohorts, respectively. Error bars represent 95% confidence intervals from 1,000 bootstrapping samples. **B, D, F** Number of edges in the correlation network after applying a 0.01 FDR cutoff as a function of the dataset sample size in the Korčula 2010, Split and Vis cohorts, respectively. Error bars represent 95% confidence intervals of 1,000 bootstrapping samples. Note that for parcor, correlation values can only be estimated for a sample size greater or equal to the number of variables, in this case 50.

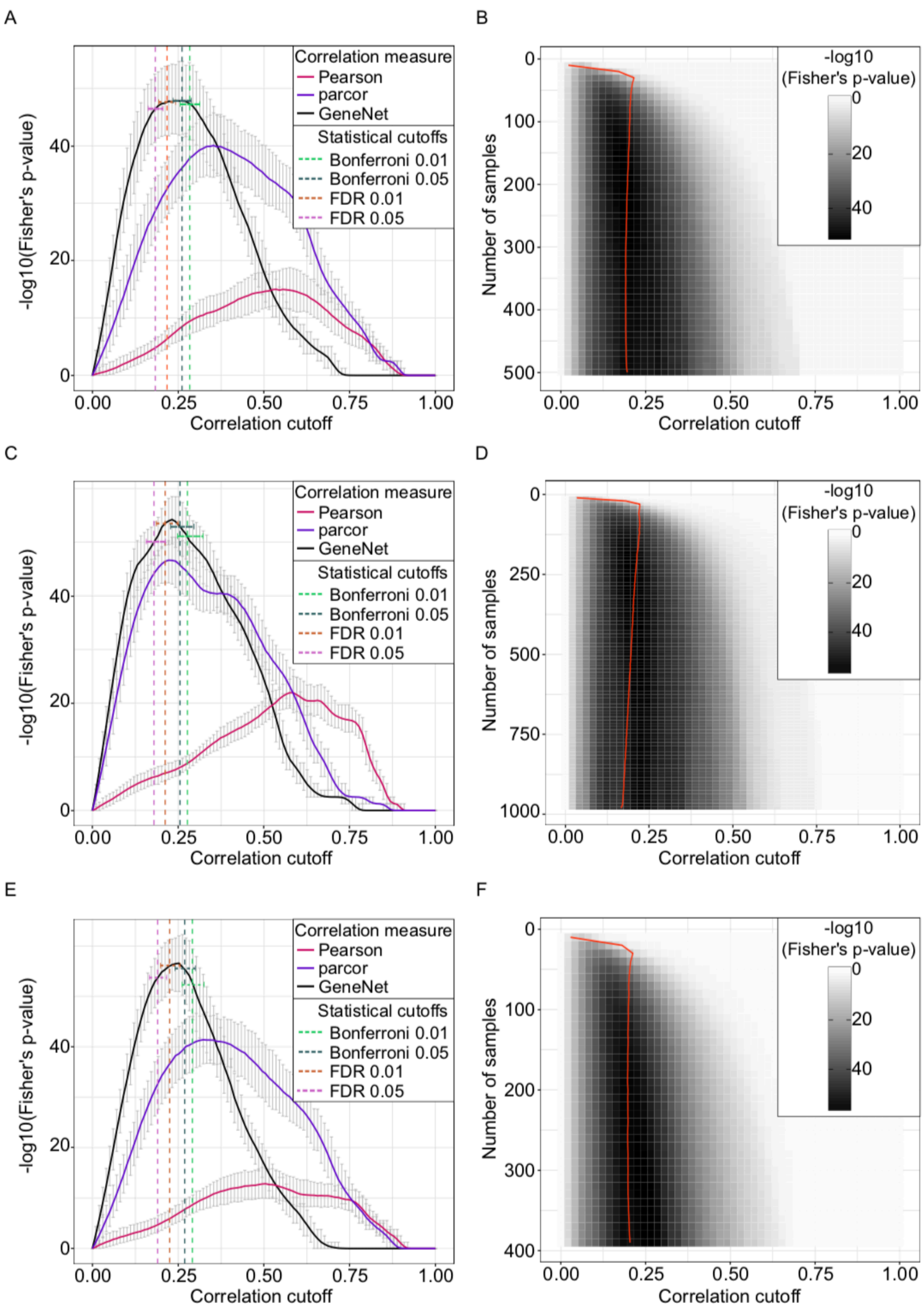


**Figure S2: Network quality as a function of the correlation cutoff in the glycomics replication cohorts.**

**A, C, E** Fisher's exact test p-values as a function of the correlation cutoff calculated in the Korčula 2010, Split and Vis cohorts, respectively, for three correlation estimators: Pearson correlation (red), exact partial correlation (purple), GeneNet partial correlation (black). For each correlation cutoff, the original dataset was bootstrapped 1,000 times. Error bars represent the 95% confidence intervals of the bootstrapping results. Dashed lines represent the mean of the bootstrapped statistical cutoffs for GeneNet. Interestingly, the minima of the GeneNet curve are fairly similar across cohorts: 0.24 (Korčula 2010), 0.23 (Split), 0.24 (Vis). **B, D, F** Fisher's exact test p-value for partial correlations estimated with GeneNet in the Korčula 2010, Split and Vis cohorts, respectively, as a function of both sample size and correlation cutoff. Colors represent the mean across 1,000 bootstrapping samples, while the red line represents the mean of the 0.01 FDR cutoff.

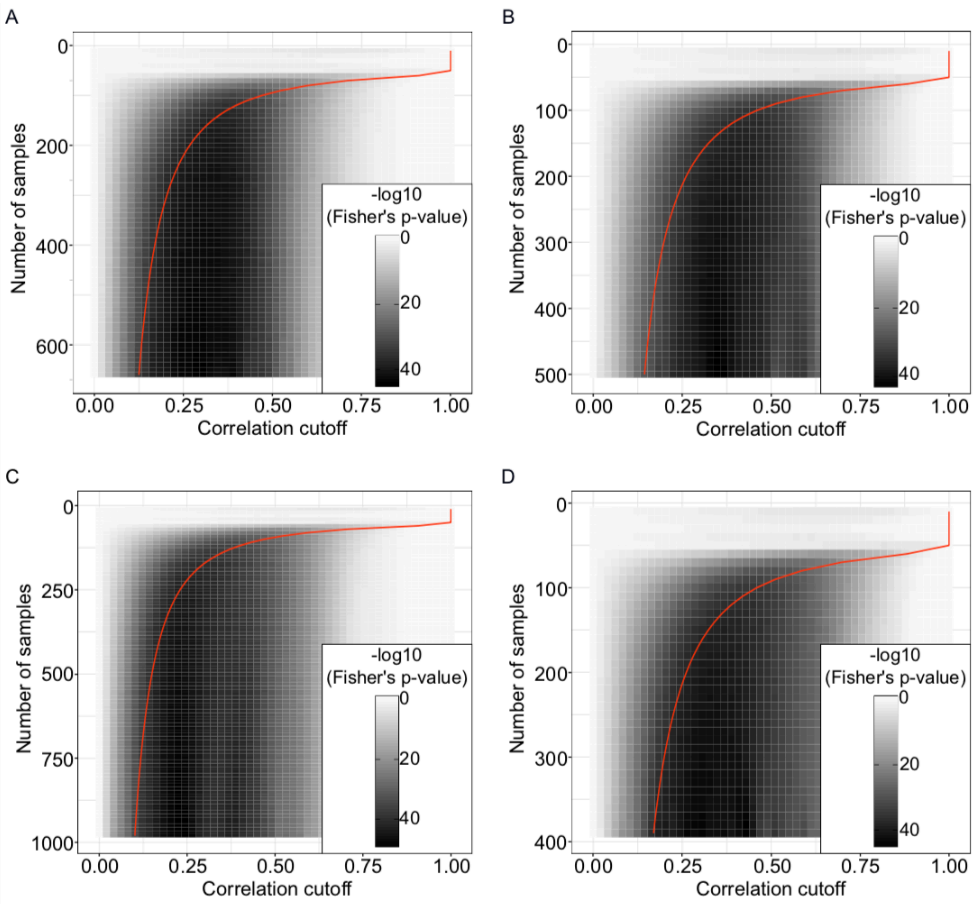


**Figure S3: Cutoff optimization as a function of the sample size for parcor correlation in the glycomics replication cohorts.**

**A, B, C, D** Fisher’s exact test p-value for partial correlations estimated with parcor in the Korčula 2013 (A), Korčula 2010 (B), Split (C) and Vis (D) cohorts, respectively, as a function of both sample size and correlation cutoff. Color represents the mean across 1,000 bootstrapping samples, while the red line represents the mean of the 0.01 FDR cutoff.

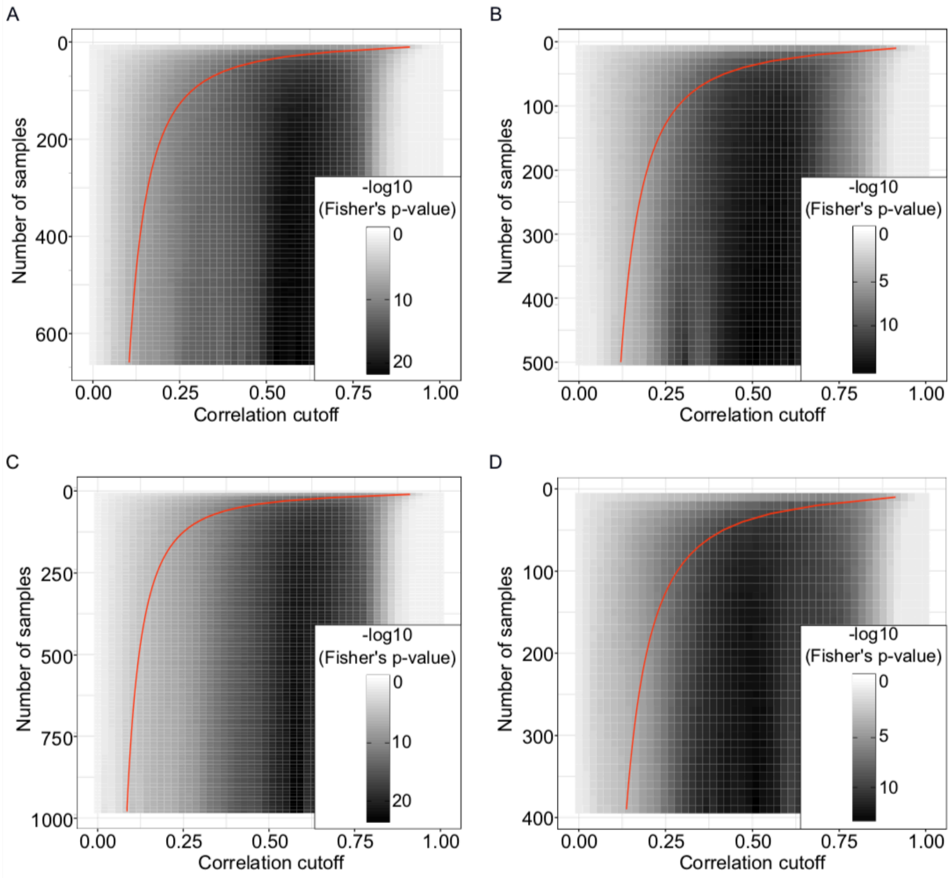


**FigureS4: Cutoff optimization as a function of the sample size for Pearson correlation in the glycomics replication cohorts.**

**A, B, C, D** Fisher's exact test p-value for Pearson correlations in the Korčula 2013 (A), Korčula 2010 (B), Split (C) and Vis (D) cohorts, respectively, as a function of both sample size and correlation cutoff. Color represents the mean across 1,000 bootstrapping samples, while the red line represents the mean of the 0.01 FDR cutoff.

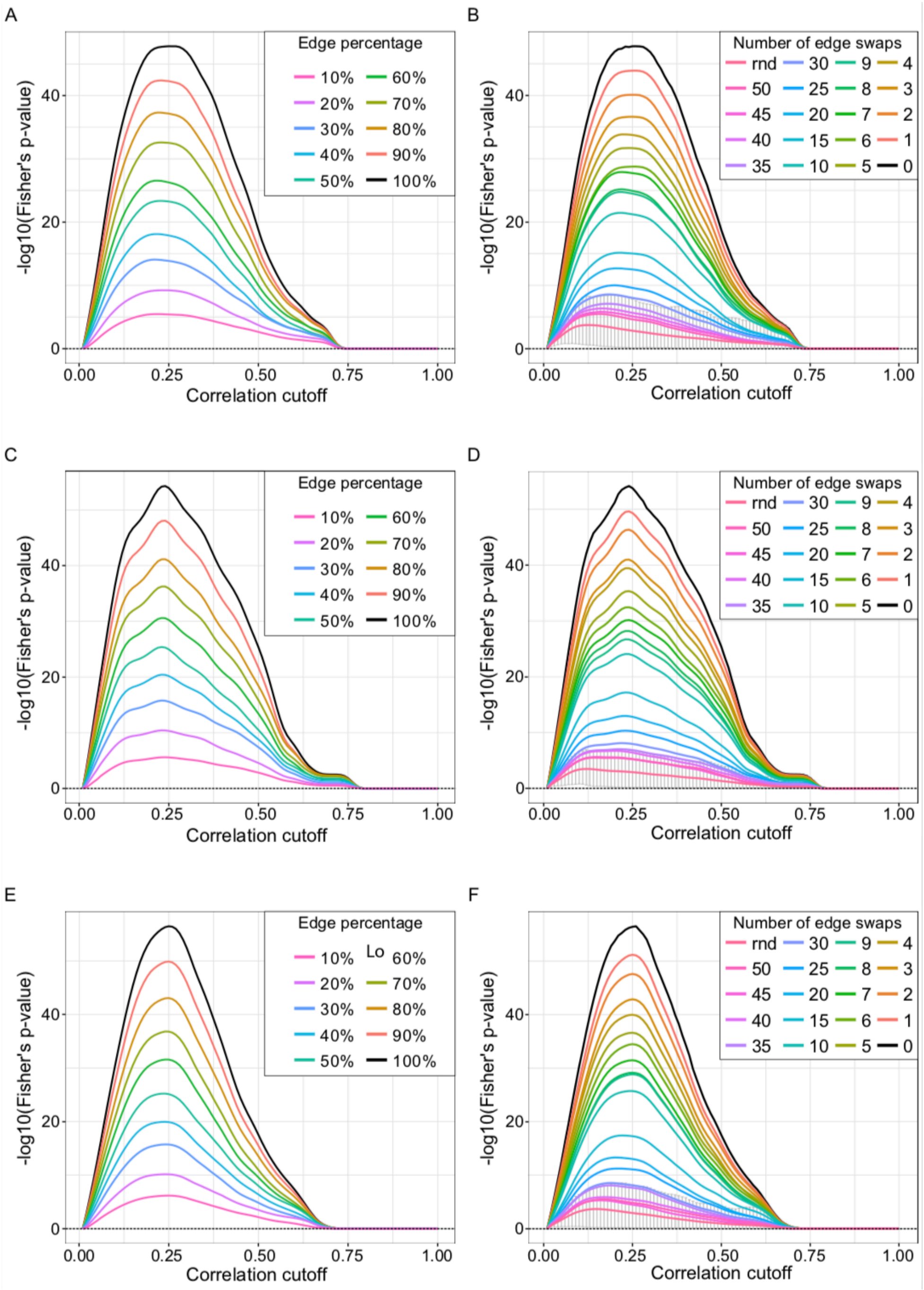


**Figure S5: Cutoff optimization with partial knowledge in the glycomics replication cohorts.**

**A, C, E** Incomplete biological reference in the Korčula 2010 (A), Split (C) and Vis (E) cohorts. For each percentage, 100 different adjacency matrices were generated by randomly selecting edges from the IgG glycosylation pathway. The curves in the figure represent the means of the 1,000 bootstrapping resamplings on each adjacency matrix. **B** Incorrect biological reference in the Korčula 2010 (B), Split (D) and Vis (F) cohorts. Edges in the IgG glycosylation pathway were randomly swapped to simulate incorrect information in the biological reference. For each considered number of swaps, 100 adjacency matrices were generated and the averages over those curves and over the 1,000 bootstrapping resamplings are shown. Here, the red curve represents 100 fully randomized adjacency matrices. The error bars on this curve represent the 95% confidence interval of the bootstrapping. Any signal that falls within these intervals should be regarded as noise.

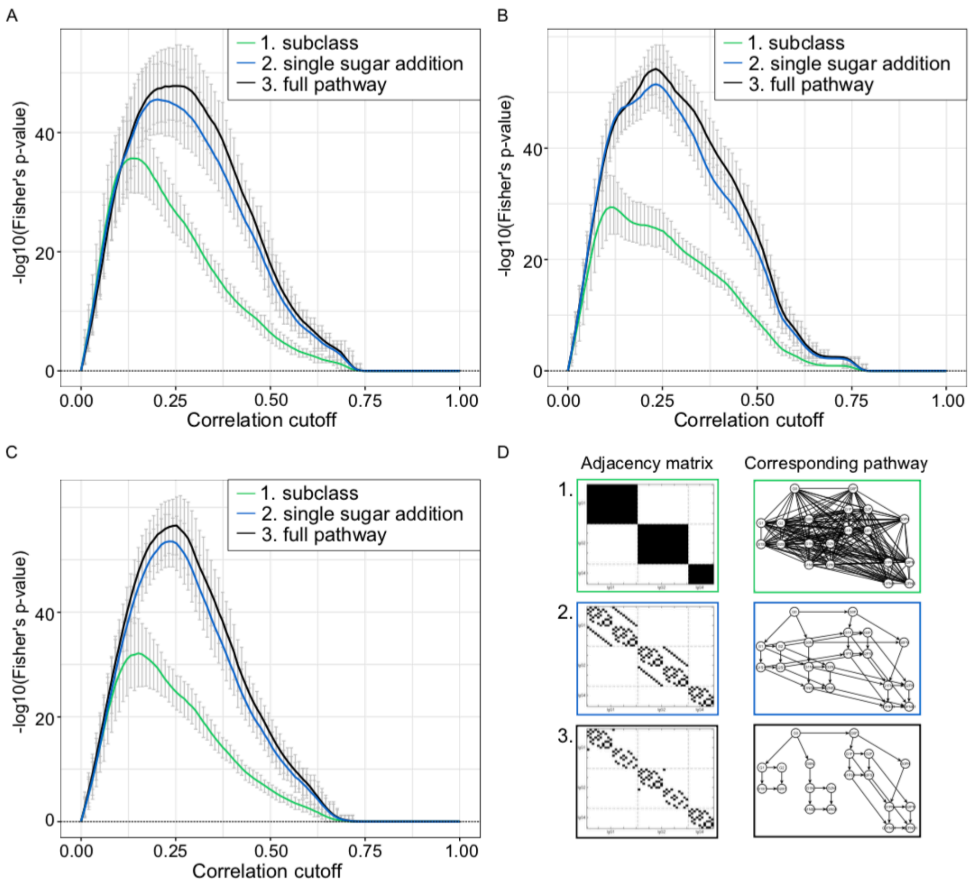


**Figure S6: Cutoff optimization with coarse prior knowledge in the glycomics replication cohorts.**

**A, B, C** Coarse biological reference in the Korčula 2010 (A), Split (B) and Vis (C) cohorts. The curves in the figure represent the means of the 1,000 bootstrapping resamplings and the different considered adjacencies. The black curve corresponds to the optimization performed on the full reference for comparison. The error bars represent the 95% confidence interval of the bootstrapping. **D** References considered in the optimization. For IgG glycomics data we know that only enzymatic reactions between glycans attached to the same IgG isoform are feasible (adjacency matrix 1) and, in addition, that only they can be modified by the addition of one sugar unit at a time (adjacency matrix 2). The black curve corresponds to the optimization performed on the full reference (adjacency matrix 3) for comparison.

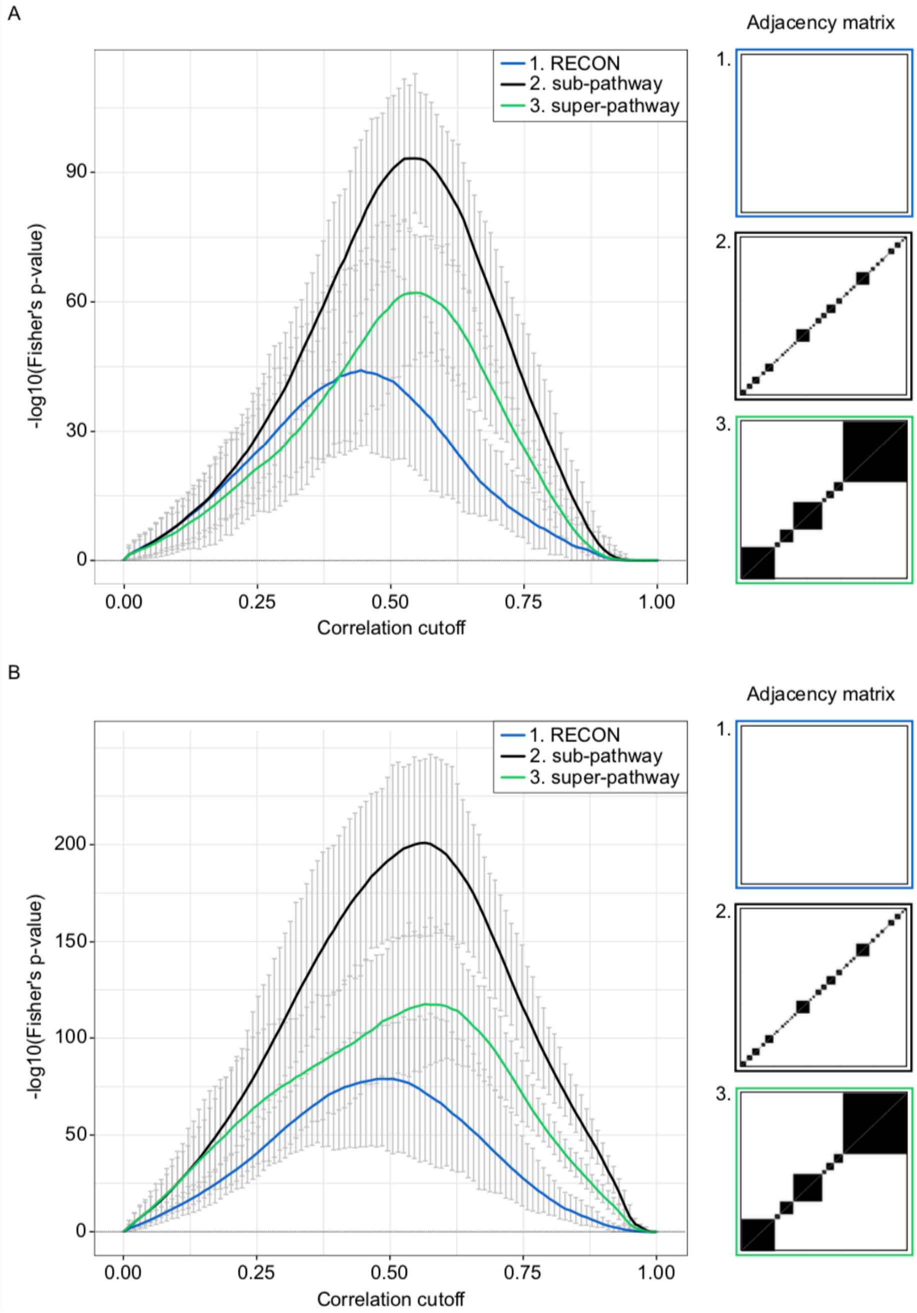


**Figure S7: Parcor (A) and Pearson (B) correlation cutoff optimization with partial and coarse knowledge in the metabolomics cohort.**

As prior knowledge, we used biochemical reactions from the RECON database (adjacency matrix 1), as well as sub- and super-pathway annotations (adjacency matrices 2 and 3, respectively). Curves in the figure represent the average over 100 bootstrapping resamplings, and error bars show the corresponding 95% confidence intervals. Vertical lines indicate the mean of the statistical cutoffs, and the horizontal error bars the corresponding 95% confidence intervals over the bootstrapping.

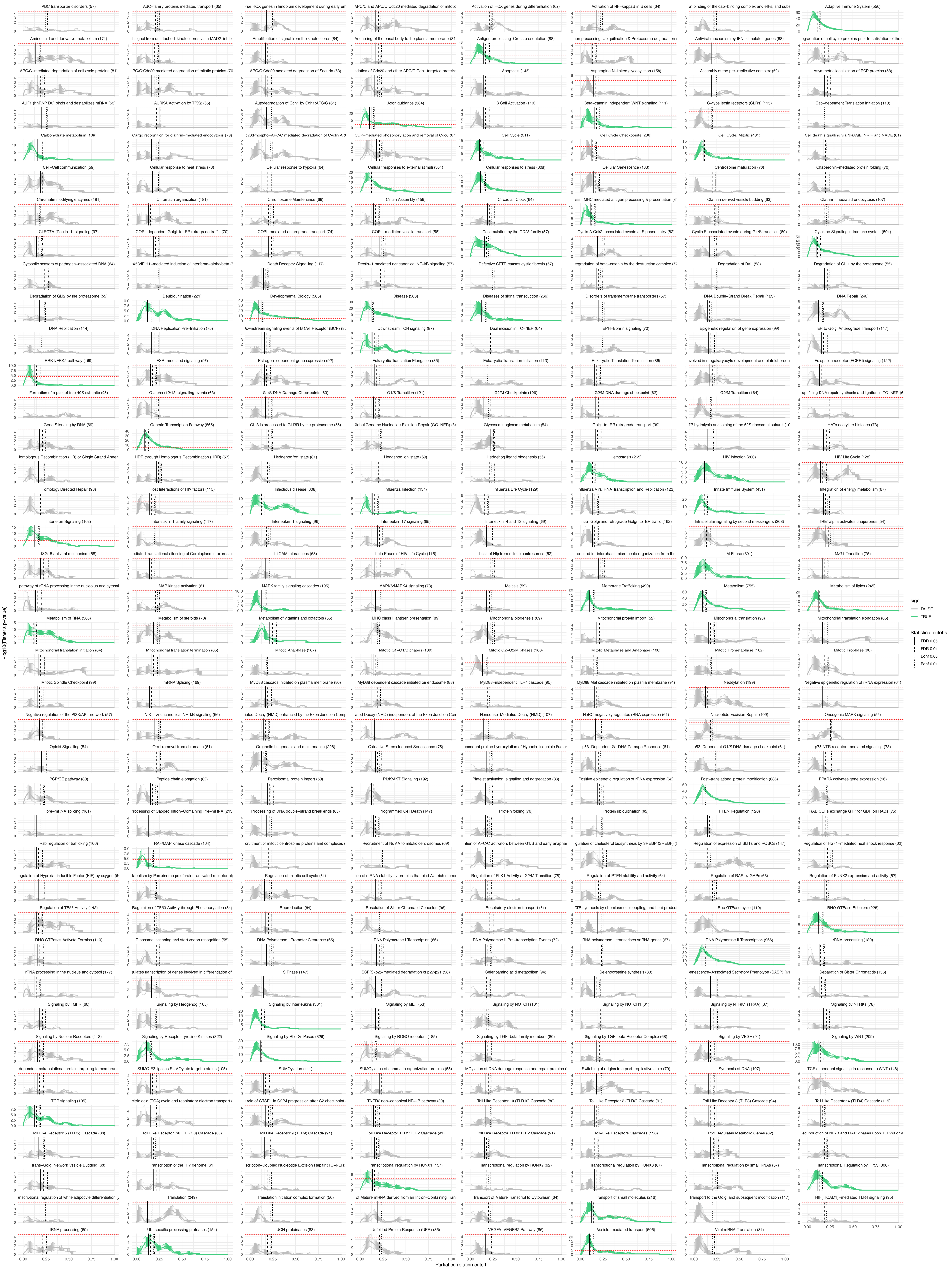
ABC transporter disorders (57)
ABC−family proteins mediated transport (65)
rior HOX genes in hindbrain development during early em
APC/C and APC/C:Cdc20 mediated degradation of mitotic
Activation of HOX genes during differentiation (62)
Activation of NF−kappaB in B cells (64)
on binding of the cap−binding complex and eIFs, and subs
Adaptive Immune System (556)
Amino acid and derivative metabolism (171)
of signal from unattached kinetochores via a MAD2 inhibi
Amplification of signal from the kinetochores (84)
Anchoring of the basal body to the plasma membrane (84)
Antigen processing−Cross presentation (88)
en processing: Ubiquitination & Proteasome degradation
Antiviral mechanism by IFN−stimulated genes (68)
egradation of cell cycle proteins prior to satisfation of the c
APC/C−mediated degradation of cell cycle proteins (81)
APC/C:Cdc20 mediated degradation of mitotic proteins (70)
APC/C:Cdc20 mediated degradation of Securin (63)
adation of Cdc20 and other APC/C:Cdh1 targeted proteins
Apoptosis (145)
Asparagine N−linked glycosylation (158)
Assembly of the pre−replicative complex (59)
Asymmetric localization of PCP proteins (58)
AUF1 (hnRNP D0) binds and destabilizes mRNA (53)
AURKA Activation by TPX2 (65)
Autodegradation of Cdh1 by Cdh1:APC/C (61)
Axon guidance (384)
B Cell Activation (110)
Beta−catenin independent WNT signaling (111)
C−type lectin receptors (CLRs) (115)
Cap−dependent Translation Initiation (113)
Carbohydrate metabolism (109)
Cargo recognition for clathrin−mediated endocytosis (73)
lc20:Phospho−APC/C mediated degradation of Cyclin A (
CDK−mediated phosphorylation and removal of Cdc6 (67)
Cell Cycle (511)
Cell Cycle Checkpoints (236)
Cell Cycle, Mitotic (431)
Cell death signalling via NRAGE, NRIF and NADE (61)
Cell−Cell communication (59)
Cellular response to heat stress (78)
Cellular response to hypoxia (64)
Cellular responses to external stimuli (354)
Cellular responses to stress (308)
Cellular Senescence (133)
Centrosome maturation (70)
Chaperonin−mediated protein folding (70)
Chromatin modifying enzymes (181)
Chromatin organization (181)
Chromosome Maintenance (69)
Cilium Assembly (159)
Circadian Clock (64)
ss I MHC mediated antigen processing & presentation (3
Clathrin derived vesicle budding (63)
Clathrin−mediated endocytosis (107)
CLEC7A (Dectin−1) signaling (97)
COPI−dependent Golgi−to−ER retrograde traffic (70)
COPI−mediated anterograde transport (74)
COPII−mediated vesicle transport (58)
Costimulation by the CD28 family (57)
Cyclin A:Cdk2−associated events at S phase entry (82)
Cyclin E associated events during G1/S transition (80)
Cytokine Signaling in Immune system (501)
Cytosolic sensors of pathogen−associated DNA (64)
DX58/IFIH1−mediated induction of interferon−alpha/beta (
Death Receptor Signalling (117)
Dectin−1 mediated noncanonical NF−kB signaling (57)
Defective CFTR causes cystic fibrosis (57)
egradation of beta−catenin by the destruction complex (7
Degradation of DVL (53)
Degradation of GLI1 by the proteasome (55)
Degradation of GLI2 by the proteasome (55)
Deubiquitination (221)
Developmental Biology (565)
Disease (563)
Diseases of signal transduction (266)
Disorders of transmembrane transporters (57)
DNA Double−Strand Break Repair (123)
DNA Repair (246)
DNA Replication (114)
DNA Replication Pre−Initiation (75)
ownstream signaling events of B Cell Receptor (BCR) (8
Downstream TCR signaling (87)
Dual incision in TC−NER (64)
EPH−Ephrin signaling (70)
Epigenetic regulation of gene expression (99)
ER to Golgi Anterograde Transport (117)
ERK1/ERK2 pathway (169)
ESR−mediated signaling (97)
Estrogen−dependent gene expression (92)
Eukaryotic Translation Elongation (85)
Eukaryotic Translation Initiation (113)
Eukaryotic Translation Termination (86)
nvolved in megakaryocyte development and platelet produ
Fc epsilon receptor (FCERI) signaling (122)
Formation of a pool of free 40S subunits (95)
G alpha (12/13) signalling events (63)
G1/S DNA Damage Checkpoints (63)
G1/S Transition (121)
G2/M Checkpoints (126)
G2/M DNA damage checkpoint (62)
G2/M Transition (164)
ap−filling DNA repair synthesis and ligation in TC−NER (6
Gene Silencing by RNA (69)
Generic Transcription Pathway (865)
GLI3 is processed to GLI3R by the proteasome (55)
lobal Genome Nucleotide Excision Repair (GG−NER) (84
Glycosaminoglycan metabolism (54)
Golgi−to−ER retrograde transport (99)
TP hydrolysis and joining of the 60S ribosomal subunit (10
HATs acetylate histones (73)
omologous Recombination (HR) or Single Strand Anneal
HDR through Homologous Recombination (HRR) (57)
Hedgehog 'off' state (81)
Hedgehog 'on' state (69)
Hedgehog ligand biogenesis (56)
Hemostasis (265)
HIV Infection (200)
HIV Life Cycle (128)
Homology Directed Repair (98)
Host Interactions of HIV factors (115)
Infectious disease (308)
Influenza Infection (134)
Influenza Life Cycle (129)
Influenza Viral RNA Transcription and Replication (123)
Innate Immune System (431)
Integration of energy metabolism (67)
Interferon Signaling (162)
Interleukin−1 family signaling (117)
Interleukin−1 signaling (96)
Interleukin−17 signaling (65)
Interleukin−4 and 13 signaling (69)
Intra−Golgi and retrograde Golgi−to−ER traffic (162)
Intracellular signaling by second messengers (208)
IRE1alpha activates chaperones (54)
ISG15 antiviral mechanism (68)
ediated translational silencing of Ceruloplasmin expressic
L1CAM interactions (63)
Late Phase of HIV Life Cycle (115)
Loss of Nlp from mitotic centrosomes (62)
required for interphase microtubule organization from the
M Phase (301)
M/G1 Transition (75)
pathway of rRNA processing in the nucleolus and cytosol
MAP kinase activation (61)
MAPK family signaling cascades (195)
MAPK6/MAPK4 signaling (73)
Meiosis (59)
Membrane Trafficking (490)
Metabolism (755)
Metabolism of lipids (245)
Metabolism of RNA (566)
Metabolism of steroids (70)
Metabolism of vitamins and cofactors (55)
MHC class II antigen presentation (89)
Mitochondrial biogenesis (69)
Mitochondrial protein import (52)
Mitochondrial translation (90)
Mitochondrial translation elongation (85)
Mitochondrial translation initiation (84)
Mitochondrial translation termination (85)
Mitotic Anaphase (167)
Mitotic G1−G1/S phases (139)
Mitotic G2−G2/M phases (166)
Mitotic Metaphase and Anaphase (168)
Mitotic Prometaphase (162)
Mitotic Prophase (90)
Mitotic Spindle Checkpoint (99)
mRNA Splicing (169)
MyD88 cascade initiated on plasma membrane (80)
MyD88 dependent cascade initiated on endosome (88)
MyD88−independent TLR4 cascade (95)
MyD88:Mal cascade initiated on plasma membrane (91)
Neddylation (199)
Negative epigenetic regulation of rRNA expression (64)
Negative regulation of the PI3K/AKT network (57)
NIK−−>noncanonical NF−kB signaling (56)
iated Decay (NMD) enhanced by the Exon Junction Comp
ated Decay (NMD) independent of the Exon Junction Com
Nonsense−Mediated Decay (NMD) (107)
NoRC negatively regulates rRNA expression (61)
Nucleotide Excision Repair (109)
Oncogenic MAPK signaling (55)
Opioid Signalling (54)
Orc1 removal from chromatin (61)
Organelle biogenesis and maintenance (228)
Oxidative Stress Induced Senescence (75)
pendent proline hydroxylation of Hypoxia−inducible Factor
p53−Dependent G1 DNA Damage Response (61)
p53−Dependent G1/S DNA damage checkpoint (61)
p75 NTR receptor−mediated signalling (78)
PCP/CE pathway (80)
Peptide chain elongation (82)
Peroxisomal protein import (53)
PI3K/AKT Signaling (192)
Platelet activation, signaling and aggregation (83)
Positive epigenetic regulation of rRNA expression (62)
Post−translational protein modification (886)
PPARA activates gene expression (96)
pre−mRNA splicing (161)
Processing of Capped Intron−Containing Pre−mRNA (213
Processing of DNA double−strand break ends (65)
Programmed Cell Death (147)
Protein folding (76)
Protein ubiquitination (65)
PTEN Regulation (120)
RAB GEFs exchange GTP for GDP on RABs (75)
Rab regulation of trafficking (106)
RAF/MAP kinase cascade (164)
cruitment of mitotic centrosome proteins and complexes (
Recruitment of NuMA to mitotic centrosomes (69)
tion of APC/C activators between G1/S and early anaphas
gulation of cholesterol biosynthesis by SREBP (SREBF) (
Regulation of expression of SLITs and ROBOs (147)
Regulation of HSF1−mediated heat shock response (62)
egulation of Hypoxia−inducible Factor (HIF) by oxygen (6
tabolism by Peroxisome proliferator−activated receptor alp
Regulation of mitotic cell cycle (81)
ion of mRNA stability by proteins that bind AU−rich eleme
Regulation of PLK1 Activity at G2/M Transition (78)
Regulation of PTEN stability and activity (64)
Regulation of RAS by GAPs (63)
Regulation of RUNX2 expression and activity (62)
Regulation of TP53 Activity (142)
Regulation of TP53 Activity through Phosphorylation (84)
Reproduction (64)
Resolution of Sister Chromatid Cohesion (96)
Respiratory electron transport (81)
TP synthesis by chemiosmotic coupling, and heat product
Rho GTPase cycle (110)
RHO GTPase Effectors (225)
RHO GTPases Activate Formins (110)
Ribosomal scanning and start codon recognition (55)
RNA Polymerase I Promoter Clearance (65)
RNA Polymerase I Transcription (66)
RNA Polymerase II Pre−transcription Events (72)
RNA polymerase II transcribes snRNA genes (67)
RNA Polymerase II Transcription (966)
rRNA processing (180)
rRNA processing in the nucleus and cytosol (177)
gulates transcription of genes involved in differentiation of
S Phase (147)
SCF(Skp2)−mediated degradation of p27/p21 (58)
Selenoamino acid metabolism (94)
Selenocysteine synthesis (83)
enescence−Associated Secretory Phenotype (SASP) (61
Separation of Sister Chromatids (156)
Signaling by FGFR (60)
Signaling by Hedgehog (105)
Signaling by Interleukins (331)
Signaling by MET (53)
Signaling by NOTCH (101)
Signaling by NOTCH1 (61)
Signaling by NTRK1 (TRKA) (67)
Signaling by NTRKs (78)
Signaling by Nuclear Receptors (113)
Signaling by Receptor Tyrosine Kinases (322)
Signaling by Rho GTPases (326)
Signaling by ROBO receptors (185)
Signaling by TGF−beta family members (80)
Signaling by TGF−beta Receptor Complex (68)
Signaling by VEGF (91)
Signaling by WNT (209)
dependent cotranslational protein targeting to membrane
SUMO E3 ligases SUMOylate target proteins (105)
SUMOylation (111)
SUMOylation of chromatin organization proteins (55)
MOylation of DNA damage response and repair proteins (
Switching of origins to a post−replicative state (79)
Synthesis of DNA (107)
TCF dependent signaling in response to WNT (148)
TCR signaling (105)
citric acid (TCA) cycle and respiratory electron transport (
role of GTSE1 in G2/M progression after G2 checkpoint (
TNFR2 non−canonical NF−kB pathway (80)
Toll Like Receptor 10 (TLR10) Cascade (80)
Toll Like Receptor 2 (TLR2) Cascade (91)
Toll Like Receptor 3 (TLR3) Cascade (94)
Toll Like Receptor 4 (TLR4) Cascade (119)
Toll Like Receptor 5 (TLR5) Cascade (80)
Toll Like Receptor 7/8 (TLR7/8) Cascade (88)
Toll Like Receptor 9 (TLR9) Cascade (91)
Toll Like Receptor TLR1:TLR2 Cascade (91)
Toll Like Receptor TLR6:TLR2 Cascade (91)
Toll−Like Receptors Cascades (136)
TP53 Regulates Metabolic Genes (82)
ed induction of NFkB and MAP kinases upon TLR7/8 or 9
trans−Golgi Network Vesicle Budding (63)
Transcription of the HIV genome (61)
scription−Coupled Nucleotide Excision Repair (TC−NER)
Transcriptional regulation by RUNX1 (157)
Transcriptional regulation by RUNX2 (92)
Transcriptional regulation by RUNX3 (87)
Transcriptional regulation by small RNAs (57)
Transcriptional Regulation by TP53 (306)
nscriptional regulation of white adipocyte differentiation (
Translation (249)
Translation initiation complex formation (56)
of Mature mRNA derived from an Intron−Containing Trans
Transport of Mature Transcript to Cytoplasm (64)
Transport of small molecules (216)
Transport to the Golgi and subsequent modification (117)
TRIF(TICAM1)−mediated TLR4 signaling (95)
tRNA processing (69)
Ub−specific processing proteases (154)
UCH proteinases (83)
Unfolded Protein Response (UPR) (85)
VEGFA−VEGFR2 Pathway (86)
Vesicle−mediated transport (506)
Viral mRNA Translation (81)
−log10(Fisher's p−value)
0.00 0.25 0.50 0.75 1.00
Partial correlation cutoff
sign
FALSE
TRUE
Statistical cutoffs
FDR 0.05
FDR 0.01
Bonf 0.05
Bonf 0.01

### Figure S8: Cutoff optimization for the transcriptomics data.

For each of the 469 pathway considered, protein-protein interaction networks from STRING were used as reference. The black curve represents the average over 100 bootstrapping resamplings, and the error bars show the corresponding 95% confidence intervals. Vertical lines indicate the mean of the statistical cutoffs, and the areas the corresponding 95% confidence intervals over the bootstrapping. Pathways are shown in alphabetical order.
