## Supplementary File S2 with data for "A strategy to incorporate prior knowledge into correlation network cutoff selection": FileS2.Rmd

````
---
title: "Cutoff Optimization Results"
output: 
  flexdashboard::flex_dashboard:
    orientation: rows
    social: menu
runtime: shiny
---
  
```{r setup, include=FALSE}

library(shiny)
library(visNetwork)
library(igraph)
library(dplyr)
library(ggplot2)

# load optimization results and network data
load("FileS2Data.Rds")
# define cutoff vector
cut_vec <- (0:100)/100

```

Sidebar {.sidebar data-width=350}
=====================================

This page allows to explore the cutoff optimization results for all TCGA transcriptomics pathways considered.

After selecting a pathway from the drop-down menu below, the results will be automatically visualized in the two tabs ("Networks", "Optimization Curve"). 

Network tab: statistical FDR 0.05 partial correlation network (upper left), the optimized partial correlation network (upper right), the reference network used for optimization based on STRING (lower left), and the reference network used for validation based on CORUM (lower right). Black edges indicate connections found in the STRING training adjacency, green edges indicate connections found in the CORUM testing adjacency, while gray edges indicate false positives.

Optimization Curve tab: cutoff optimization results with 100 bootstrap samples. The line indicates the median of the bootstrap, and the error bars indicate the 95% confidence intervals.


```{r}

selectInput('pws', 'Pathway', sort(names(net_data)), selected = "Axon guidance")

```

Networks
===================================== 

Row 
-------------------------------------
  
### Statistical Network (FDR 0.05)
  
```{r}

renderVisNetwork(
    {
      visNetwork(nodes=net_data[[input$pws]]$nn_fdr, edges = net_data[[input$pws]]$ee_fdr) %>%
        visIgraphLayout(layout = "layout.norm", layoutMatrix = as.matrix(net_data[[input$pws]]$Coords[,1:2]))  %>%
        return
    })

```


### String-Optimized Network

```{r}

renderVisNetwork(
    {
visNetwork(nodes=net_data[[input$pws]]$nn_opt, edges = net_data[[input$pws]]$ee_opt) %>%
  visIgraphLayout(layout = "layout.norm", layoutMatrix = as.matrix(net_data[[input$pws]]$NetCoords_opt[,2:3])) %>%
        return
    })

```


Row
-------------------------------------
  
### Training Adjacency - STRING

```{r}

renderVisNetwork(
    {
visNetwork(nodes=net_data[[input$pws]]$nn_s, edges = net_data[[input$pws]]$ee_s) %>%
      visIgraphLayout(layout = "layout.norm", layoutMatrix = as.matrix(net_data[[input$pws]]$NetCoords_s[,2:3])) %>%
        return
    })

```

### Testing Adjacency - CORUM

```{r}

renderVisNetwork(
    {
visNetwork(nodes=net_data[[input$pws]]$nn_c, edges = net_data[[input$pws]]$ee_c) %>%
      visIgraphLayout(layout = "layout.norm", layoutMatrix = as.matrix(net_data[[input$pws]]$NetCoords_c[,2:3])) %>%
        return
    })

```
  
Optimization Curve
===================================== 

```{r}

data_plot <- reactive({
  as.data.frame(cbind(cut_vec=cut_vec, data[[input$pws]]$res_opt$res_adjl$s$ntw_cut_offs_ps[-c(1:4),]))
})

renderPlot({
  p <- ggplot(data_plot(), aes(x=cut_vec, y=-M))+
        geom_errorbar(aes(ymin=-U, ymax=-L),color="grey50")+
        geom_line(size=1) +
        xlab("Partial correlation cutoff")+
        ylab("-log10(Fisher's p-value)")+
        theme_light(base_size=10) +
        theme(strip.text = element_text(size=6))+
        geom_vline(aes(xintercept=m0, 
                       linetype=rownames(data[[input$pws]]$res_opt$res_adjl$s$stat_cutoffs)),
                       data=as.data.frame(data[[input$pws]]$res_opt$res_adjl$s$stat_cutoffs))+
        scale_linetype_manual(values=c(3,4,2,1),
                          name="Statistical cutoffs",
                          breaks=c("fdr_0.05","fdr_0.01","bonf_0.05","bonf_0.01"),
                          labels=c("FDR 0.05", "FDR 0.01", "Bonf 0.05", "Bonf 0.01")) +
        geom_hline(yintercept = -log10(0.01/length(data)), color ="red", linetype="dashed",size=0.5)+
        ggtitle("Cutoff optimization based on the STRING adjacency")
  
  print(p)
})

```
````
